# SLC-25A46 Regulates Mitochondrial Fusion through FZO-1/Mitofusin and is Essential for Maintaining Neuronal Morphology

**DOI:** 10.1101/2024.02.11.579862

**Authors:** Hiroyuki Obinata, Hironori Takahashi, Satoshi Shimo, Toshiyuki Oda, Asako Sugimoto, Shinsuke Niwa

**Author notes:** Corresponding author: SHINSUKE NIWA, Frontier Research Institute for Interdisciplinary Sciences (FRIS), Tohoku University, Aramaki-Aoba 6-3, Aoba-Ku, Sendai, Miyagi 980-0845, Japan.

## Abstract

Mitochondria are dynamic organelles shaped by sequential fission and fusion events. The mitochondrial protein SLC25A46 has been identified as a causative gene for mitochondrial neuropathies. However, the function of SLC25A46 in mitochondrial morphogenesis remains controversial, with several reports suggesting it acts as a mitochondrial fission factor, while others propose it as a fusion factor. In this study, employing forward genetics, we identified *slc-25A46*, a *Caenorhabditis elegans* orthologue of human SLC25A46, as an essential factor for mitochondrial fusion. Suppressor mutagenesis screening revealed loss-of-function mutations in *drp-1*, a mitochondrial fission factor, as suppressors of *slc-25A46*. The phenotype of *slc-25A46* mutants is similar to those of *fzo-1* mutants, wherein the mitochondrial fusion factor Mitofusin is disrupted. Overexpressing FZO-1/Mitofusin mitigated mitochondrial defects in *slc-25a46* mutants, indicating SLC-25A46 promotes fusion through FZO-1/Mitofusin. Disease model worms carrying mutations associated with SLC25A46 exhibited mitochondrial fragmentation and accelerated neurodegeneration, suggesting *slc-25A46* maintains neuronal morphology through mitochondrial fusion regulation.

## Introduction

Mitochondria are highly dynamic organelles that constantly change their shapes to adapt to the metabolic states and physiological environments of the cell ^1^. The morphological diversities of mitochondria arise from sequential fission and fusion activities regulated by a family of GTPase enzymes ^1^. In humans, mitochondrial fission is mediated by dynamin-related protein 1 (DRP1), a GTPase that forms ring-like structures at division sites^2,3^. Conversely, mitochondrial fusion requires other GTPases, such as Mitofusin 1 and 2 (MFN1 and 2) for outer membrane (OM) fusion and Optic atrophy 1 (OPA1) for inner membrane (IM) fusion^4–6^. In yeast, the fusion process involves the participation of another protein called Ugo1, which localizes to OM and coordinates fusion between the outer and inner membranes by interacting with Mitofusin and OPA1 orthologs ^7,8^.

A mitochondrial protein SLC25A46, originally identified as a cause of optic atrophy and spastic paraplegia, shares weak similarities with Ugo1^9^. Like Ugo1, SLC25A46 is localized on the OM, forms a complex with MFNs and OPA1^10^. Despite these similarities, the exact function of SLC25A46 in mitochondrial dynamics remains controversial. Several studies have reported that acute knockdown of SLC25A46 in cultured cell lines leads to the formation of large elongated mitochondria^9–11^. Knockdown of SLC25A46 has been shown to cause elongated mitochondria in zebrafish and Drosophila^12–14^.A similar elongated mitochondrial phenotype has been observed in SLC25A46 mutant mice that have a 46-bp deletion in the exon 8 ^15^. Conversely, overexpression of SLC25A46 induces mitochondrial fragmentation ^9^ . All of these studies support the function of SLC25A46 as a fission factor. In contrast, the complete loss of SLC25A46 in the knock-out Hela cell results in the fragmentation of the mitochondrial network^16^. In a separate study using a knockout mouse model, loss of SLC25A46 function led to the formation of small mitochondrial fragments in the nervous system^17^. A mutation in bovine SLC25A46 gene is associated with sensorimotor neuropathy, known as Turning calves syndrome, wherein fragmented aggregated mitochondria were observed^17^. Moreover, a null mutation in human SLC25A46 gene has been identified as a cause of optic atrophy spectrum disprder^18^. In patient cells with this mutation, mitochondrial fragmentation was observed^18^. These phenotypes are similar to those observed in yeast Ugo1 mutants, suggesting that primary role of SLC25A46 is as a mitochondrial fusion factor.

*Caenorhabditis elegans* is a widely-used model organism for studying organelle dynamics and morphogenesis, including the nucleus, synaptic vesicles and mitochondria ^19–24^. Disruption of intracellular transport of mitochondria is observed in *unc-116*, *miro-1* and *trak-1* mutant worms, where the molecular motor kinesin-1 and its adaptor proteins are mutated^20,21^ . Orthologues of these proteins have similar functions in mammals and flies^24^. Mitochondrial fragmentation has been reported in *fzo-1* and *eat-3* mutant worms^25^. *fzo-1* and *eat-3* respectively encode MFN1/2 and OPA1 orthologues. Inversely, mitochondrial hyperfusion has been observed in *drp-1* mutants in which a DRP1 orthologue is disrupted ^26^ ^27^.

In this study, through forward genetic screening in *C. elegans*, we identified a loss-of-function mutation in *slc-25A46* gene, a *C. elegans* orthologue of human SLC25A46. The *slc-25A46* mutant worms exhibited fragmented and small mitochondria, similar to the phenotype observed in worms lacking *fzo-1/Mitofusin*. Notably, this defect was partially rescued by a loss-of-function mutation in *drp-1* or overexpressing *fzo-1*. These findings suggest that *slc-25A46* is upstream of *fzo-1* and is essential for mitochondrial fusion, rather than fission. To further investigate the function of SLC25A46, we introduced pathogenic mutations reported in human SLC25A46 gene into *C. elegans slc-25A46* gene and examined their effects on mitochondrial morphology and distribution. Our results indicate that *slc-25A46* mutations accelerated the morphological degeneration in neurons. Collectively, these findings suggest that *slc-25A46* plays a critical role in mitochondrial morphology and neuronal maintenance.

## Results

### Isolation of *slc-25A46* mutants

It is established that mitochondria accumulate in dendrites of ciliated sensory neurons, such as olfactory neurons ^28^. To elucidate the molecular mechanisms of mitochondrial morphogenesis in sensory dendrites, we labeled mitochondria in *C. elegans* PHA neurons (Fig. 1A). GFP was fused to the N-terminal 54 amino acids of TOMM-20 protein and used as a marker to visualize mitochondria^19,29^. The PHA neuron is a highly polarized neuron characterized by a dendrite with sensory cilia, a cell body and an axon^30^. We utilized the *flp-15* promoter, which exclusively expresses in PHA neurons^31^. In the resulting strain, intense GFP fluorescence of mitochondria was observed in both the soma and dendrites (Fig 1A). In contrast, analyzing signals in the axon was challenging due to the sparse distribution of mitochondria and the autofluorescence of gut cells.

**Figure 1.**
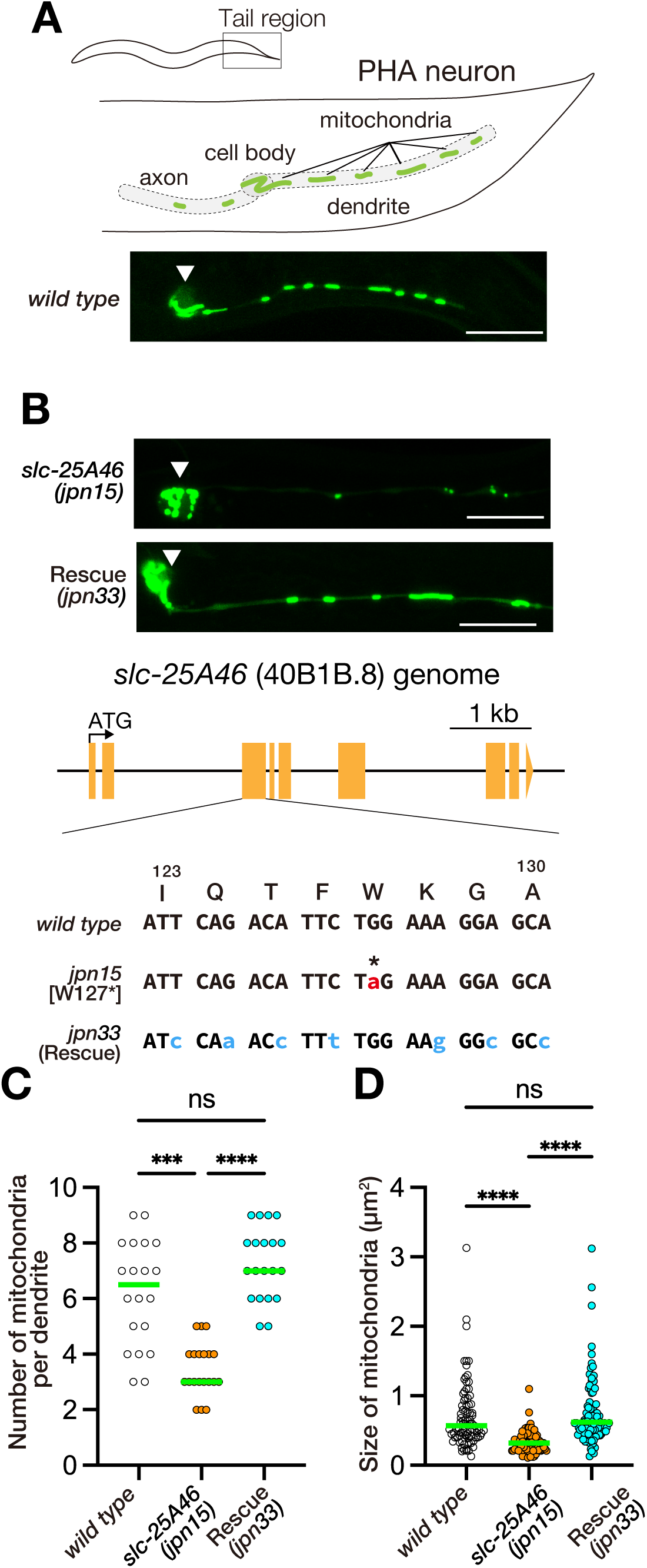
Identification of *slc-25A46* mutant. (A) Schematic of PHA neuron morphology (upper panel) and a representative image showing the morphology and distribution of mitochondria in PHA neuron (lower photo). The arrowhead indicates the cell body. TOMM-20(1-54) was fused with GFP and expressed using the *flp-15* promoter. Bar, 10 µm. (B) Alleles of *slc-25A46* and their impact on mitochondrial morphology. The *jpn15* allele, isolated through EMS mutagenesis, has a nonsense mutation in the *slc-25A46* gene and exhibits fragmented mitochondria phenotype in the PHA neuron. Using CRISPR/cas9 technology, the nonsense mutation in the *slc-25A46(jpn15)* allele was corrected. The resultant allele *slc-25A46(jpn33)* has normal mitochondria. Bars, 10 µm. (C) Dot plots showing the number of mitochondria in the PHA dendrite. Each dot shows the number of mitochondria in a single PHA dendrite. Green bars represent median values. n = 20 dendrites from 20 worms. Kruskal-Wallis test followed by Dunn’s multiple comparison test. ***, p < 0.001. ****, p < 0.0001. ns, p > 0.05 and statistically not significant. (D) Dot plots showing the size distribution of mitochondria in the PHA dendrite. Each dot represents the size of an individual mitochondrion in the PHA dendrite. Green bars represent median values. n = 85, 83 and 85 mitochondria. Kruskal-Wallis test followed by Dunn’s multiple comparison test. ***, p < 0.001. ****, p < 0.0001. ns, p > 0.05 and statistically not significant.

To identify molecules essential for mitochondrial morphogenesis, we conducted forward genetic screens through genome-wide ethyl methanesulfonate (EMS) mutagenesis^32^. We isolated a mutant allele named *jpn15*. In *jpn15* worms, the number of mitochondria was reduced, and their size was smaller compared to wild type (Fig 1B). Through genetic mapping and genome sequencing, we identified a nonsense mutation at W127 of the *slc-25A46* gene, which is the *C. elegans* orthologue of the mammalian SLC25A46 (Fig 1B). Previous RNAi screen of mitochondrial morphology have not linked the function of s*lc-25A46* with mitochondrial morphogenesis ^19^. To confirm that W127Stop mutation in the *slc-25A46* gene is the causative mutation for the *jpn15* phenotype, we used CRISPR/Cas9-mediated genome editing to substitute the nonsense mutation in *jpn15* with the *wild-type* amino acid. We introduced silent mutations and created a restriction enzyme site in the repair template to prevent subsequent Cas9 cleavage and verify successful genome editing (Fig 1B). The resultant allele, *jpn33*, carrying the repaired *slc-25A46*, showed restored mitochondrial number and size, indicating that the W127Stop mutation in *slc-25A46* locus is responsible for the abnormal mitochondrial morphology observed in *jpn15* allele (Fig 1B-D). Moreover, we found that *slc-25A46(gk570223)*, obtained from the million mutation project^33^, had a Q310Stop mutation and showed the mitochondrial fragmentation phenotype (Supplementary Figure S1A).

Next, to determine if mitochondrial fragmentation occurs in other tissues of *slc-25A46(jpn15)* mutant worms, we observed mitochondria within the body wall muscles. Similar to the observations in the PHA neuron cell bodies, mitochondria in *slc-25A46(jpn15)* mutants exhibited rounded and fragmented structures compared to wild type (Supplementary Figure S1B). Prior studies have shown that cristae structures are degenerated by SLC25A46 mutations in human and mice ^10,15,16^, whereas a study has shown that cristae structures are not strongly affected in SLC25A46-knockout mice^17^. We observed the fine structure of mitochondria in the muscular cells by transmission electron microscopy. Our observation suggested that cristae structures in mitochondria was not strongly affected in *slc-25A46* mutant worms (Supplementary Figure S1C and D), resembling the findings described in Duchesne et al.(2017) ^17^.

Overall, our findings suggest that *slc-25A46* plays a crucial role in maintaining the size and distribution of mitochondria.

### *drp-1* mutation suppress mitochondrial fragmentation in *slc-25A46* mutants

To understand the function of SLC25A46 in mitochondrial morphogenesis, we performed second EMS mutagenesis screens and searched for mutations that can suppress the mitochondrial fragmentation phenotype in the *slc-25A46(jpn15)* mutant. A mutant allele, named *jpn73*, could recover the mitochondrial network in the cell body of *slc-25A46(jpn15)* mutant (Fig 2A). Mapping and sequencing revealed that *jpn73* is an allele of *drp-1* mutant. Genome sequencing revealed that exon 1 of the *drp-1* gene, including the start codon, was deleted in *drp-1(jpn73)* allele (Fig 2B). DRP-1 is a *C. elegans* orthologue of DRP1 that is essential for the mitochondrial fission^26,27^. We next compared the phenotype of *drp-1(jpn73)* and *slc-25A46(jpn15)* single mutants (Fig 2C). Moreover, we observed *drp-1(tm1108)*, which is a null allele of *drp-1*, for comparison (Fig 2B). In the dendrite of PHA neuron, both *drp-1(jpn73)* and *drp-1(tm1108)* mutants had an elongated mitochondrion, indicating mitochondrial hyperfusion (Fig 2C). The phenotype of *drp-1* was clearly different from mitochondrial fragmentation phenotype observed in *slc-25A46(jpn15)* (Fig 1 and 2C). These results shows that SLC-25A46 and DRP-1 have opposite functions in the mitochondrial morphogenesis. As DRP-1 is a mitochondrial fission factor, it is suggested that SLC-25A46 works as a mitochondrial fusion factor.

**Figure 2.**
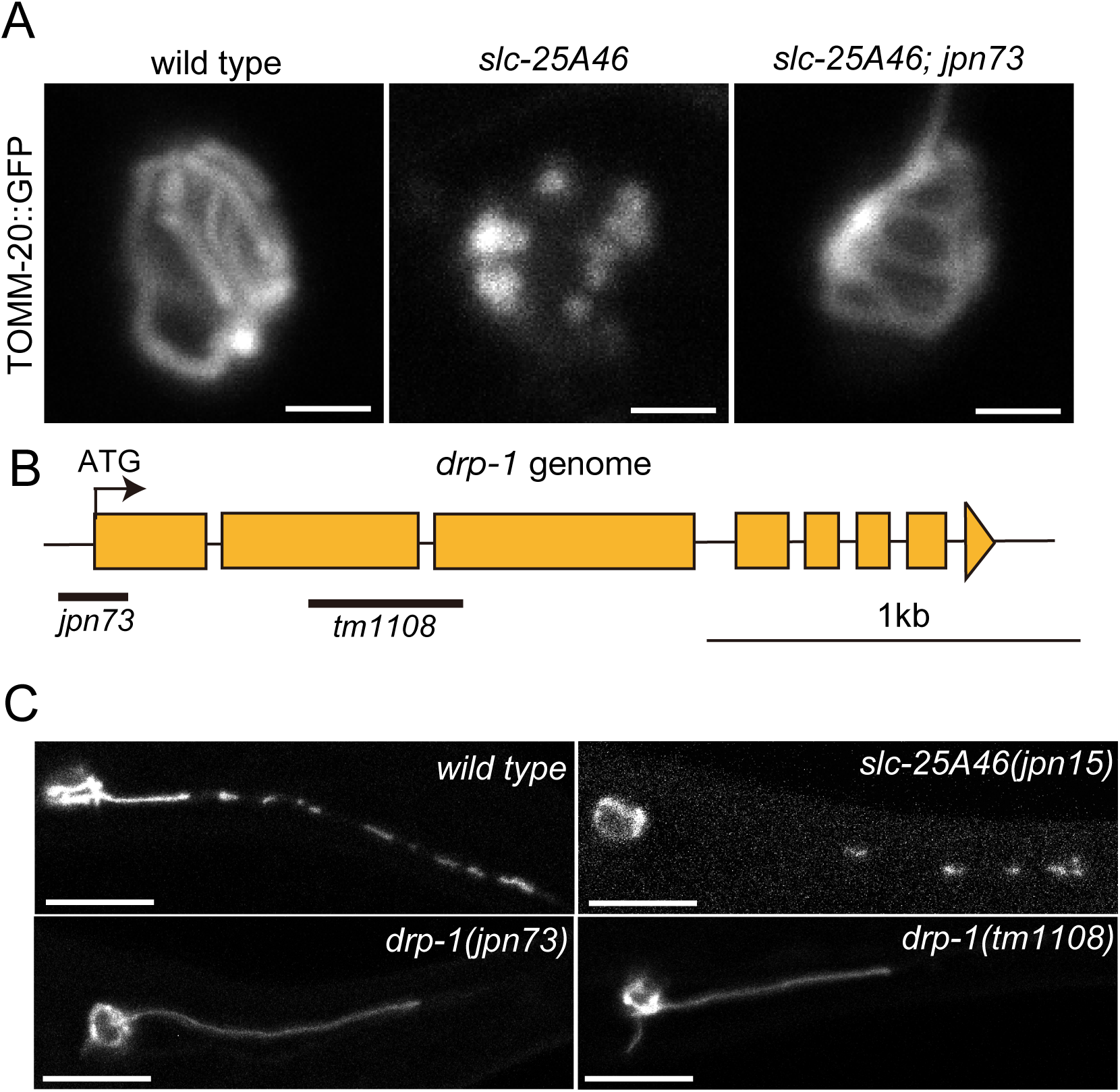
SLC-25A46 and DRP1 have opposite functions in mitochondrial morphogenesis. (A) Identification of *jpn73*, a suppressor mutant of the *slc-25A46* mutant. Representative images showing mitochondria in the cell body of *wild type*, *slc-25A46*, *slc-25A46*; *jpn73* mutants. Note that fragmented mitochondria phenotype in *slc-25A46* is suppressed in *slc-25A46; jpn73* double mutant. Bars, 2 µm. (B) Schematic drawing showing the genome structure of *drp-1* gene and the *jpn73* mutation. Deleted regions in *jpn73* and *tm1108* alleles are shown by bars. (C) Representative images showing the mitochondrial morphology in the dendrite of *wild type*, *slc-25A46(jpn15)*, *drp-1(jpn73)* and *drp-1(tm1108)*. Bars, 10 µm.

### *slc-25A46* works as a mitochondrial fusion factor in *C. elegans*

To study the phenotype of *slc-25A46*, we performed a comparative analysis of mitochondrial morphology using *C. elegans* mutants with loss-of-function mutations in key genes associated with mitochondrial dynamics, including *drp-1* (Fig 2), *fzo-1* and *eat-3* (Fig 3A). FZO-1 and EAT-3, orthologues of MFN1/2 and OPA1 respectively, play important roles in mitochondrial fusion (Fig 3A). We firstly focused on the morphology of mitochondria within the cell bodies of PHA neurons. Mitochondria were categorized into tubular, fragmented, and aggregated forms (Fig 3B). Within the fragmented category, a further distinction was made between weak and strong fragmentation based on the size of the fragmented mitochondria (Fig 3B). In the cell body of *wild type* and *drp-1(tm1108)* mutants, we could find a tubular mitochondrial network (Fig 3C and D). No fragmented mitochondria were observed in *wild type* and *drp-1(tm1108)* mutants. In contrast, the cell body of the *slc-25A46 (jpn15)* mutant contained fragmented and small mitochondria, similar to cell bodies of *fzo-1(tm1133)* and *eat-3(tm1107)* mutants (Fig 3C and D). However, the degree of fragmentation in *slc-25A46(jpn15)* was milder than that in *fzo-1(tm1133)* and *eat-3(tm1107)* mutants (Fig 3D). Next, we compared the number and morphology of mitochondria in the PHA dendrite. In *slc-25A46(jpn15)* mutants, there was a significant decrease in both the number and size of mitochondria compared to the *wild type* (Figs 3E-G). Similarly, *fzo-1(tm1133)* mutants displayed reductions in both the number and size of mitochondria. Interestingly, *eat-3(tm1107)* mutants only exhibited a decrease in mitochondrial size, without abnormal localization. Based on these phenotypes, it is suggested that SLC-25A46 acts as a fusion factor for mitochondria, similar to FZO-1 and EAT-3. Notably, the phenotype of *slc-25A46* is more similar to that of *fzo-1* than to that of *eat-3*.

**Figure 3.**
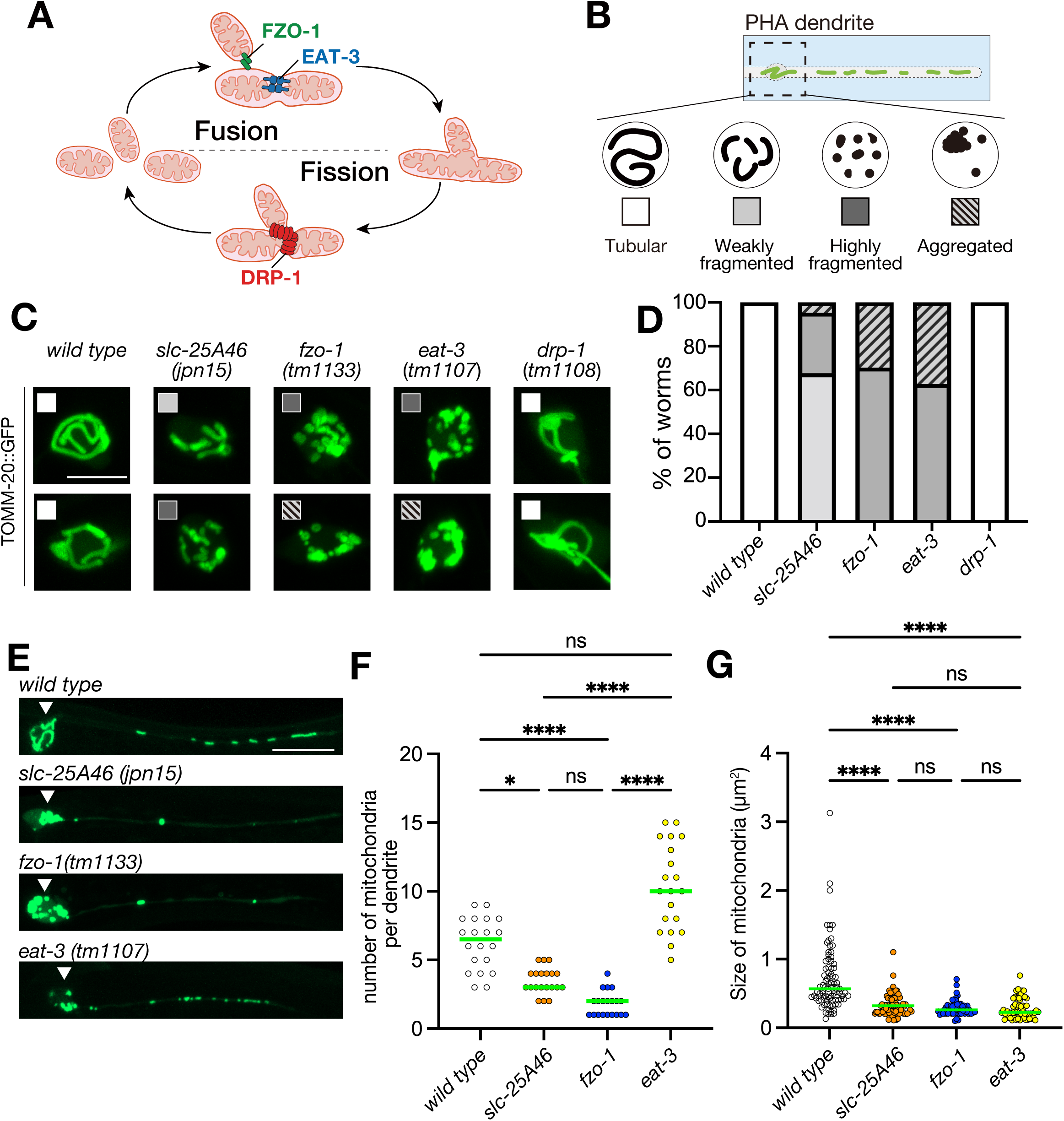
(A) Schematic drawing showing the mitochondrial dynamics and functions of FZO-1(Mitofusin), EAT-3(OPA1) and DRP-1(DRP1). (B) Schematic drawing showing the category of mitochondrial morphology used in this study. (C and D) Mitochondrial morphology in the cell body of PHA neuron. Representative images showing the mitochondrial morphology in PHA cell body and their categories (C). In panel (D), a stacked bar graph illustrates the ratio of categories in each mutant. N = 40 worms. Bars, 2 µm. (E) Representative images showing mitochondrial morphology in the PHA neuron. Bars, 10 µm. (F) Dot plots showing the number of mitochondria in the PHA dendrite. Each dot shows the number of mitochondria in a single PHA dendrite. Green bars represent median values. n = 20 dendrites from 20 worms. Kruskal-Wallis test followed by Dunn’s multiple comparison test. *, p < 0.05. ****, p < 0.0001. ns, p > 0.05 and statistically not significant. (G) Dot plots showing the size distribution of mitochondria in the PHA dendrite. Each dot represents the size of an individual mitochondrion in the PHA dendrite. Green bars represent median values. n = 85, 83, 53 and 60 mitochondria for each genotype. Kruskal-Wallis test followed by Dunn’s multiple comparison test. ****, p < 0.0001. ns, p > 0.05 and statistically not significant.

### Ectopic Expression of FZO-1 partially suppresses defects in *slc-25a46* mutant

The mammalian SLC25A46 has been reported to physically interact with the outer and inner membrane fusion factors MFN1 and MFN2 (FZO-1 orthologs) ^11^. The above genetic experiments suggested that *slc-25A46* and *fzo-1* work together in mitochondrial morphogenesis. However, the functional interaction between SLC25A46 and MFNs remains elusive. To investigate the relation further, we conducted additional genetic experiments. Firstly, we generated *slc-25A46*; *fzo-1* double mutants and found that the phenotype was not enhanced compared with either *slc-25A46* or *fzo-1* single mutants (Supplementary Figure S2). We next expressed FZO-1 in the *slc-25A46* mutant (Fig 4). As a result, the ectopic expression of FZO-1::mCherry in PHA neurons was able to restore both the number and size of mitochondria in *slc-25A46 (jpn15)* mutant (Fig 4A-C). These results collectively suggest that *slc-25A46* is functionally upstream of *fzo-1* in the regulation of mitochondrial morphology.

**Figure 4.**
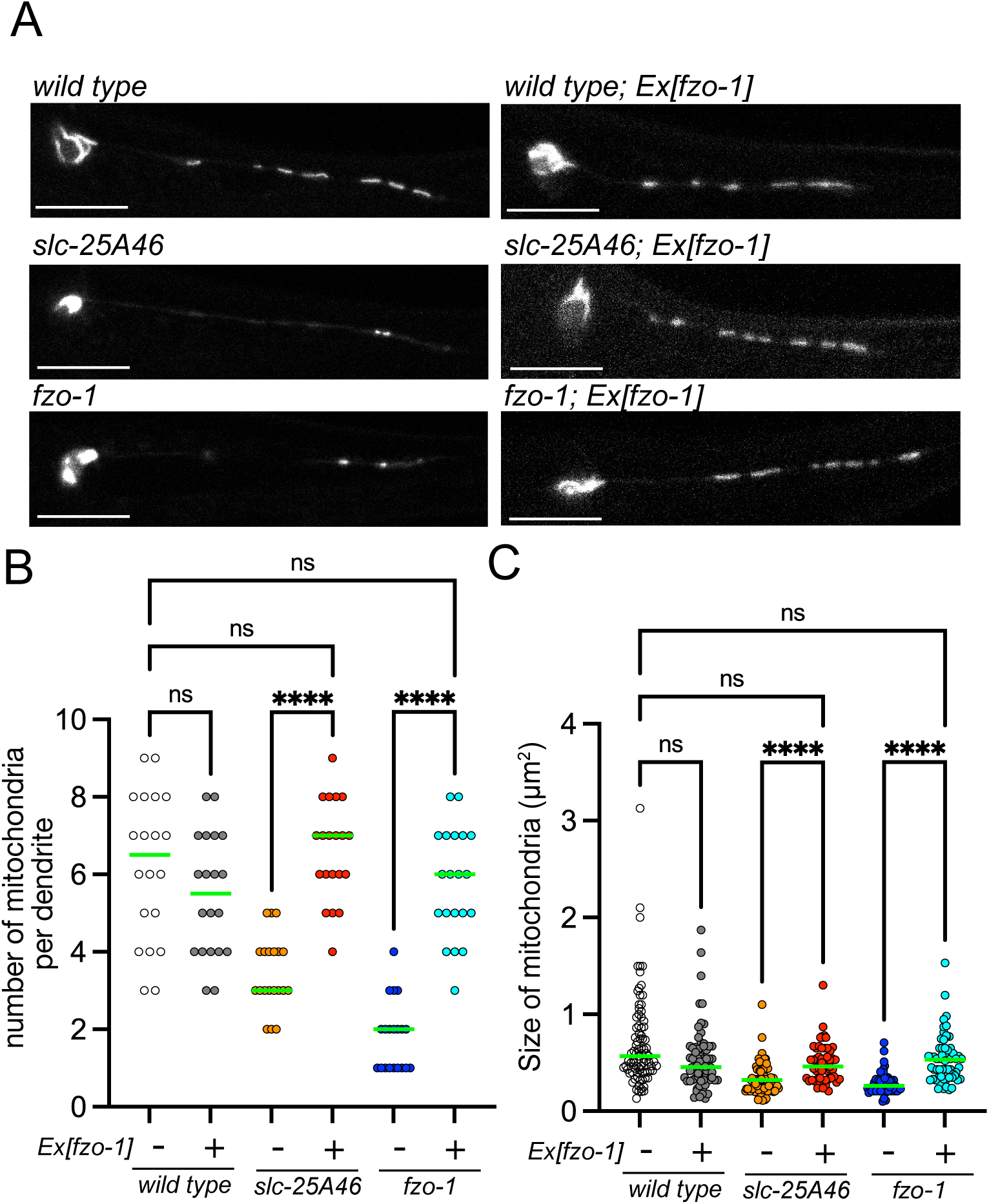
Ectopic expression of FZO-1 suppresses the mitochondrial morphology in the slc-25A46 mutant. (A) Representative images showing the mitochondrial morphology in the PHA neuron of *wild type*, *slc-25A46(jpn15)* and *fzo-1* mutants, as well as strains overexpressing FZO-1 in each mutant background. Note that overexpression of FZO-1 rescue mitochondrial defects in both *slc-25A46(jpn15)* and *fzo-1* mutants. Bars, 5 µm. (B) Dot plots showing the number of mitochondria in the PHA dendrite. Each dot shows the number of mitochondria in a single PHA dendrite. Green bars represent median values. n = 20 dendrites from 20 worms. Kruskal-Wallis test followed by Dunn’s multiple comparison test. ****, p < 0.0001. ns, p > 0.05 and statistically not significant. (C) Dot plots showing the size distribution of mitochondria in the PHA dendrite. Each dot represents the size of an individual mitochondrion in the PHA dendrite. Green bars represent median values. n = 85, 70, 83, 57, 53 and 60 mitochondria, respectively. Kruskal-Wallis test followed by Dunn’s multiple comparison test. ****, p < 0.0001. ns, p > 0.05 and statistically not significant.

### Ectopic Expression of SLC-25A46 induce mitochondrial fragmentation

Previous studies have shown that overexpression of SLC25A46 induces mitochondrial fragmentation in vertebrate systems ^9^. Thus, we overexpressed SLC-25A46 in *C. elegans* and observed the mitochondrial morphology (Fig 5A-C). Similar to vertebrate systems, the overexpression of SLC-25A46 induced mitochondrial fragmentation in the *wild-type* background (Fig 5A-C). Moreover, we found that the expression of SLC-25A46 did not rescue fragmented mitochondria phenotype in either the *slc-25A46* or *fzo-1* mutant background. These results suggest that the proper expression level of SLC-25A46 is essential for mitochondrial fusion, as discussed later.

**Figure 5.**
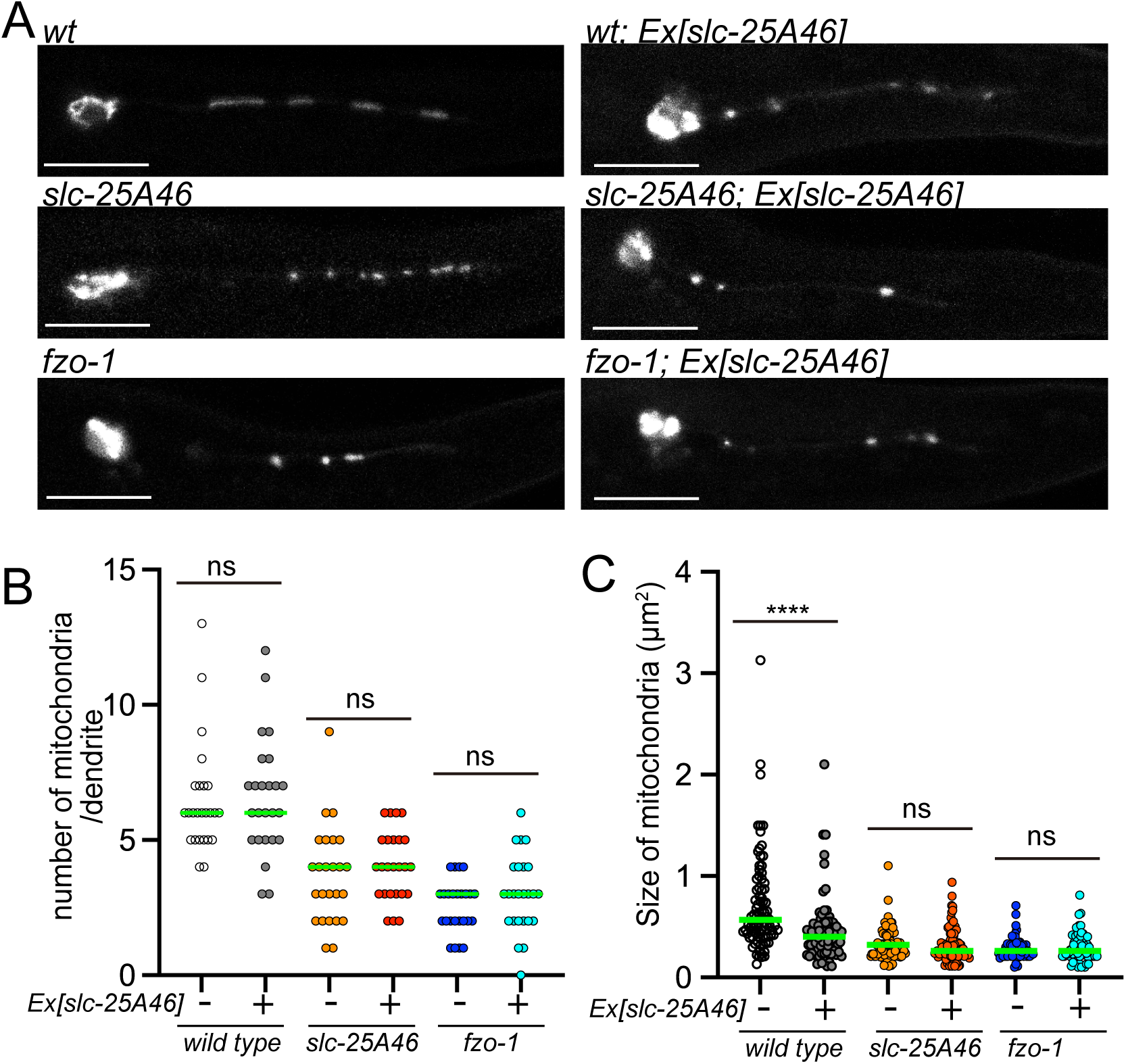
Ectopic expression of SLC-25A46 induces fragmentation of mitochondria. (A) Representative images showing the mitochondrial morphology in the PHA neuron of *wild type*, *slc-25A46(jpn15)* and *fzo-1* mutants, as well as strains overexpressing SLC-25A46 in each mutant background. Note that overexpression of SLC-25A46 caused fragmentation of mitochondria in the wild type but overexpression of SLC-25A46 could not rescue mitochondrial defects in *slc-25A46(jpn15)* and *fzo-1* mutants. Bars, 5 µm. (B) Dot plots showing the number of mitochondria in the PHA dendrite. Each dot shows the number of mitochondria in a single PHA dendrite. Green bars represent median values. n = 20 dendrites from 20 worms. Kruskal-Wallis test followed by Dunn’s multiple comparison test. ns, p > 0.05 and statistically not significant. (C) Dot plots showing the size distribution of mitochondria in the PHA dendrite. Each dot represents the size of an individual mitochondrion in the PHA dendrite. Green bars represent median values. n = 85, 81, 83, 82, 53 and 64 mitochondria, respectively. Kruskal-Wallis test followed by Dunn’s multiple comparison test. ****, p < 0.0001. ns, p > 0.05 and statistically not significant.

### SLC25A46-associated disease model worms

Human SLC25A46 has been identified as a causative gene for neurodegenerative disorders such as Charcot-Marie-Tooth disease and optic atrophy ^9^. Most of these diseases are caused by autosomal recessive mutations, and numerous amino acid changes have been reported as disease-causing in humans. However, the mechanisms underlying disease onset remain unclear. Therefore, we attempted to validate the effects of disease-causing mutations on mitochondrial morphology and distribution by introducing these mutations into *C. elegans slc-25A46* gene. Among the human disease-associated mutations reported to date, we introduced three mutations into *C. elegans slc-25A46* gene by CRISPR/cas9 (Fig 6A). These residues are embedded in the OM (Fig 6B). Firstly, we observed the morphology of mitochondria in the cell body of mutant alleles. We found mitochondria are fragmented in disease-associated *slc-25A46* mutant worms (Fig 6C and D). However, compared with *slc-25A46(jpn15)*, which is considered to be a null allele, these three mutants showed more subtle mitochondrial fragmentation in the cell body (Fig. 6C and D). Next, we observed the distribution and morphology of mitochondria in the PHA dendrite (Fig 6E-G). The size of mitochondria was statistically smaller in two out of three disease mutant model worms (Fig 6F). While we could not detect statistical significance, *slc-25A46(E301D)* also tended to have smaller mitochondria. In contrast, the number of mitochondria was not significantly affected by these disease-associated mutations (Fig 6G). These data suggest that disease-associated SLC25A46 mutations result in partial loss of function in the regulation of mitochondrial fusion event.

**Figure 6.**
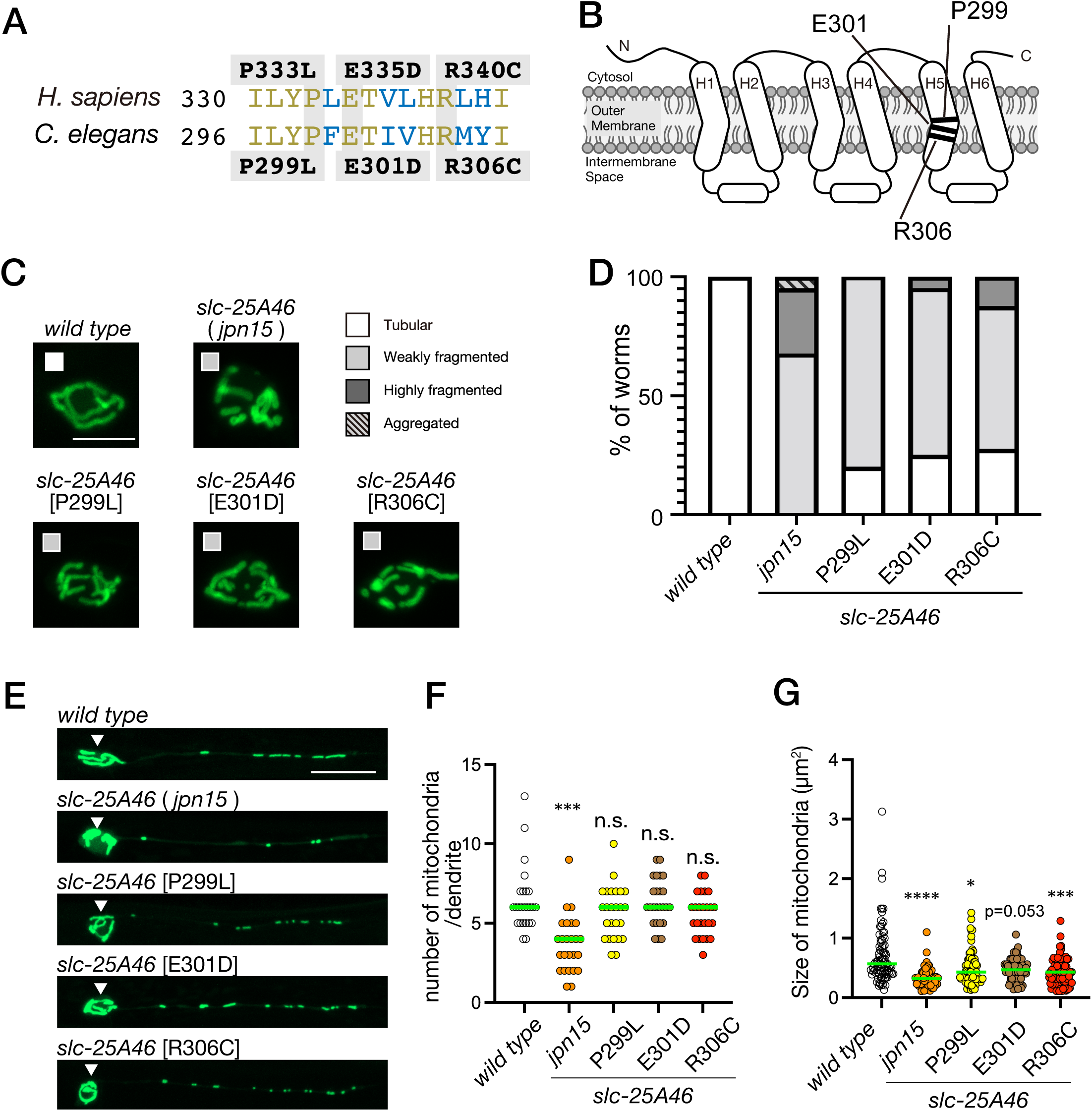
Establishment of SLC25A46-associated disease models. (A) A sequence comparison between human SLC25A46 and *C. elegans* SLC-25A46. Three amino acid changes associated with SLC25A46-related diseases (P333L, E335D and R340C) and the corresponding amino acid changes in *C. elegans*, analyzed in this study, are illustrated. (B) Schematic drawing showing the predicted structure of *C. elegans* SLC-25A46 and mutations analyzed in this study. (C and D) Mitochondrial morphology in the cell body of PHA neuron. Representative images showing the mitochondrial morphology in PHA cell body in *slc-25A46* alleles and their categories (C). In panel (D), a stacked bar graph illustrates the ratio of categories in each mutant. Mitochondrial morphology was categorized using the criteria shown in Figure 3C. N = 40 worms. Bars, 2 µm. (E) Representative images showing the mitochondrial morphology in the PHA neuron of *wild type*, *slc-25A46* mutant alleles. Arrow heads indicate the cell body. Bars, 5 µm. (F) Dot plots showing the number of mitochondria in the PHA dendrite. Each dot shows the number of mitochondria in a single PHA dendrite. Green bars represent median values. n = 20 dendrites from 20 worms for each genotype. Kruskal-Wallis test followed by Dunn’s multiple comparison test. ***, p < 0.001. ns, p > 0.05 and statistically not significant. (G) Dot plots showing the size distribution of mitochondria in the PHA dendrite. Each dot represents the size of an individual mitochondrion in the PHA dendrite. Green bars represent median values. n = 85, 83, 83, 80, 80 and 80 mitochondria for each genotype, respectively. Kruskal-Wallis test followed by Dunn’s multiple comparison test. *, p < 0.05. ***, p < 0.001. ****, p < 0.0001. Actual p value is shown for *slc-25A46(E301D)* mutant.

### The neuronal morphology is disrupted by *slc25-a46* mutations

Mutations in the human SLC25A46 gene have been linked to neuronal degeneration ^9,10^ . In light of this, we investigated the neuronal morphology in our *slc-25A46* mutant worms. Specifically, we observed the morphology of PVD neurons, a model neuronal cell often used to study neurodegeneration and regeneration in *C. elegans* ^21,34,35^ (Fig 7A). We observed adult stage worms 1 day to 7 days after the final molting. In 1-day-old adult worms, the overall morphology of the PVD neuron appeared unaffected by *slc-25A46* mutations, preserving the characteristic menorah structure ^36^ (Fig 7A and B). However, we noticed an increased number of bead-like structures along the dendrite, a reported sign of neuronal degeneration ^21^ ^37^ ^34^, in *slc-25A46* mutant worms at day 1 and day 3 (Fig 7B and 7C). At 5-day-old and 7-day-old adult worms, because the number of bead-like structures is increased even in wild-type worms, no significant difference was detected between *wild type* and *slc-25A46* mutants in the number of bead-like structures (Fig 7C). However, the morphological defects were more evident in *slc-25A46* mutant worms at day 5, showing an increased ectopic branches (Fig 7B). As a result, the area occupied by neurites was significantly increased in 5-day-old *slc-25A46* mutants compared to age-matched wild type worms (Fig 7D). Previous studies have shown that ectopic branches of PVD neurons increase when neurons are injured and failed to be repaired ^34^ ^35^. These data indicate that *slc-25A46* mutations accelerate morphological degeneration in neurons.

**Figure 7.**
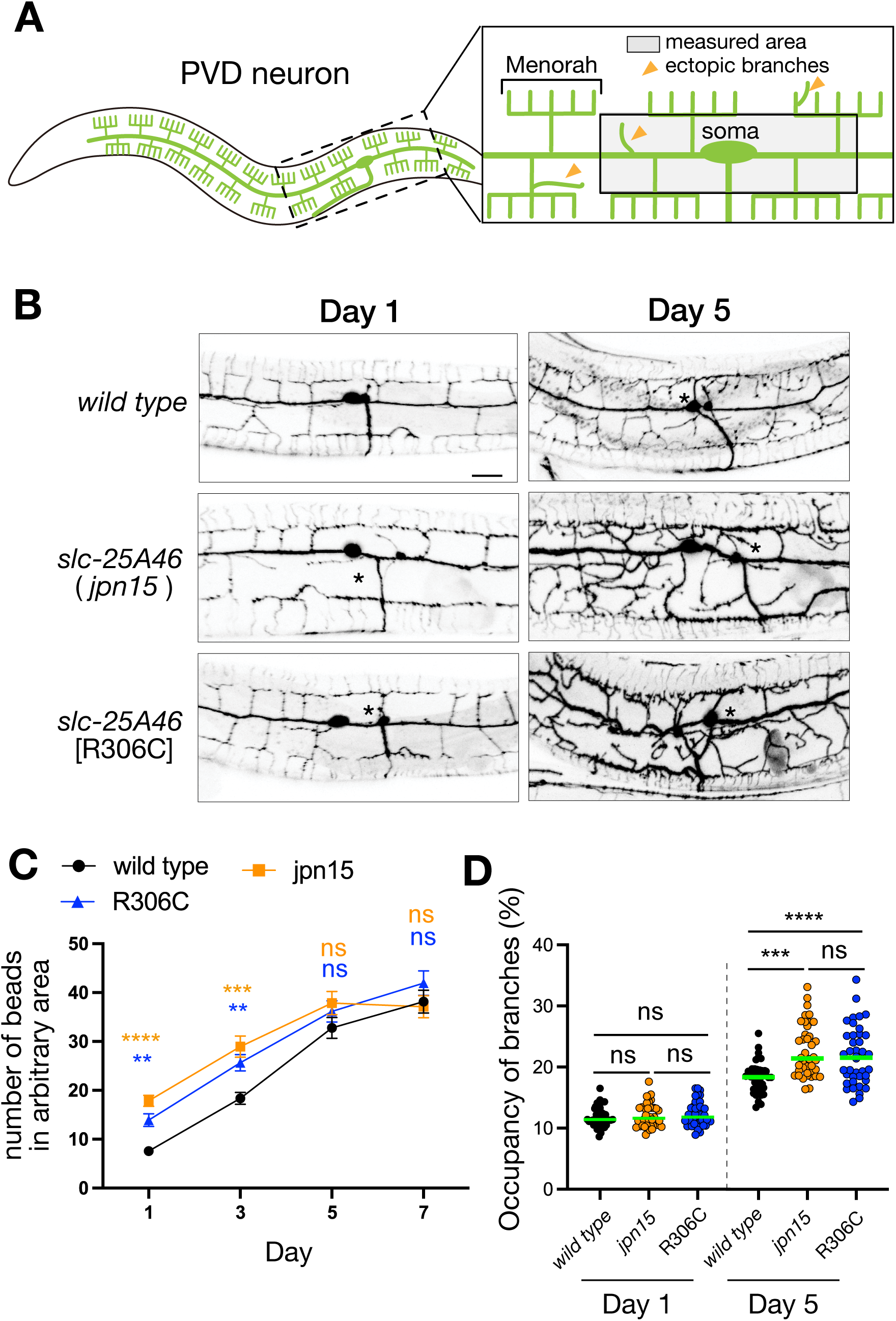
Neuronal morphology in slc-25A46 mutant worms. (A) Schematic drawing of the morphology of PVD neuron. The drawing also illustrates examples of ectopic branches. Measured area in panels C and D are shown by a gray box. (B) Representative images showing the morphology of PVD neurons in *wild type* and *slc-25A46* mutant alleles at Day 1 and 5. (C) A graph showing the number of bead-like structures along the dendrite of PVD neuron at days 1, 3, 5 and 7. Measured area is indicated in panel A. Mean ± standard errors of the mean (SEM) is shown in the graph. n = 20 worms for each genotype and each day. Kruskal-Wallis test followed by Dunn’s multiple comparison test was performed using wild type as a control. **, p < 0.01. ***, p < 0.001. ****, p < 0.0001. ns, p > 0.05 and statistically not significant. (D) Dot plots showing the percentage of area occupied by PVD neurons within the region illustrated in panel A. An elevated value is considered to represent an increase in ectopic branches. n = 40 PVD neurons from 40 worms for each genotype and each day. Kruskal-Wallis test followed by Dunn’s multiple comparison test. ***, p < 0.001. ****, p < 0.0001. ns, p > 0.05 and statistically not significant.

## Discussion

### SLC-25A46 is essential for mitochondrial fusion

The role of SLC25A46 in multicellular organisms remains controversial. Hyperfusion of mitochondria has been reported in knockdown zebrafish and Drosophila models ^14^. In mammalian systems, both fragmentation and hyperfusion of mitochondria have been reported when the expression level of SLC25A46 is perturbed. Most works reporting hyperfusion of mitochondria tend to be based on the knockdown experiments or an observation in mutant mice considered to be hypomorph, whereas studies reporting fragmentation of mitochondria tend to be based on the complete knockout. Thus, it is possible that the manipulation methods or the expression level of SLC25A46 may cause these differences. We show mitochondrial fragmentation in our *slc-25A46(jpn15)* allele which has an early stop codon and considered to be an amorph allele. We found that the mitochondrial phenotype of *slc-25A46* mutants is similar to those of *fzo-1* and *eat-3* mutants. FZO-1 is an ortholog of MFN1 and 2 whereas EAT-3 is an ortholog of OPA1, both are essential factors for mitochondrial fusion. Consistent with the role of SLC-25A46 in mitochondrial fusion, suppressor mutagenesis screening on *slc-25A46* mutant background shows that a *drp-1* mutation, which disrupt the function of mitochondrial fission factor, can antagonize the phenotype of *slc-25A46*. These genetic data suggest that *slc-25A46* is required for the mitochondrial fusion, rather than fission. On the other hand, it has been reported that fragmentation of mitochondria is induced by the overexpression of SLC25A46 in mammalian cells and zebrafish^9^. Consistent with these observations, we found overexpression of SLC-25A46 on *wild-type* background induces mitochondrial fragmentation (Fig 5). These phenomena appears to indicate that SLC-25A46 promotes mitochondrial fission. However, we have found, on the other hand, that the phenotypes of *slc-25A46* mutants cannot be rescued by the overexpression of SLC-25A46 (Fig 5). These data would show that proper expression levels of SLC-25A46 are essential for the maintenance of mitochondrial morphology, rather than indicate SLC-25A46 is a factor essential for the mitochondrial fission.

### SLC-25A46 regulates mitochondrial fusion through FZO-1

Previous studies have shown physical interactions between SLC25A46 and MFN1 and 2 ^10^. However, the functional relation between SLC25A46 and MFNs has been elusive. Taking advantage of worm genetics, we show that mitochondrial fusion mediated by *slc-25A46* is accomplished through the function of FZO-1, a worm MFN. The phenotype of *slc-25A46*; *fzo-1* double mutants are similar to those of single mutants, suggesting that SLC-25A46 and FZO-1 work in the same pathway in the mitochondrial fusion process. Moreover, overexpression of FZO-1 could partially suppress the phenotype of *slc-25A46*, indicating that SLC-25A46 is genetically upstream of FZO-1. It has been shown that FZO-1 is the GTPase that directly induces mitochondrial membrane fusion. Our genetic data suggests that SLC-25A46 helps the activity of FZO-1 in the mitochondrial fusion process. It would be interesting to biochemically test whether SLC-25A46 can enhance the GTPase activity of FZO-1.

### Worm models for mitochondria-associated disorders

The mitochondrial phenotypes in model worms suggest that disease mutations induce weak loss of function effects on the function of SLC-25A46. This is consistent with the genotype of patients; most SLC25A46 mutations cause neurological disorders in an autosomal recessive manner^9^. On the other hand, mitochondrial hyperfusion has been reported in patient cells ^9^. Further analysis is needed to determine the extent to which disease worm models reflect human symptoms and whether our model worms can be used to search for treatment methods for mitochondrial diseases. In addition to mitochondrial defects, we found acceleration of neurodegeneration in *slc-25A46* mutant worms. While we have not noticed morphological defects in the PHA neuron, we found morphological defects in the PVD neuron which is a widely used model to observe neurodegeneration in C. elegans. It has been shown that PVD neurons show morphological defects in *mtx-1* mutant worm in which mitochondrial transport is reduced^21^. Because PVD neuron is the largest neuron in *C. elegans* and has more complex dendrite structures, the morphology of PVD neuron may be more sensitive to mitochondrial defects than PHA neuron.

### Limitation of this study

While worm genetics suggest that SLC-25A46 is a mitochondrial fusion factor, we cannot exclude a possibility that worm SLC-25A46 and mammalian SLC25A46 have different functions in the mitochondrial morphogenesis. To show that the relation between SLC-25A46 and FZO-1 is conserved in mammalian cells, experiments similar to those done in this study, such as rescue of SLC25A46-knockout cells by the expression of MFN1 and 2 or analyzing of SLC25A46- and DRP1-double knockout cells, need to be repeated in mammalian systems. For that purpose, using complete knockout cells (i.e. null mutant), rather than knockdown cells or partial knockout cells (i.e. hypomorphic mutant), is required.

## Acknowledgement

We would like to thank members of Niwa lab and Sugimoto lab for helpful discussions. Some worm strains were obtained from Shen lab (Stanford University). tm alleles were obtained from NBRP. Some strains were provided by the CGC, which is funded by NIH Office of Research Infrastructure Programs (P40 OD010440). SN was supported by JSPS KAKENHI (grants nos. 22H05523 and 23H02472).

## Statement

During the preparation of this work the authors used ChatGPT in order to check English grammar and improve English writing. After using this tool, the authors reviewed and edited the content as needed and take full responsibility for the content of the publication.

## Experimental Procedures

### Plasmid preparation

Plasmids used in this study are detailed in Supplementary Table S1. Restriction enzymes were purchased from New England Biolabs (Ipswich, MA, USA). Plasmids encoding *odr-1p::gfp* and *tomm-20(1-54aa)::gfp* were obtained from Kang Shen lab. The *flp-15* promoter (*flp-15p*), a PHA neuron-specific promoter, was previously described ^31^. The DNA fragment encoding *flp-15p* was inserted between the SphI and AscI restriction enzyme sites of the *tomm-20(1-54)::gfp* plasmid. The plasmid encoding *flp-15p::slc-25A46::(GGGGS)3::mCherry* was constructed based on the plasmid encoding *flp-15p::tomm-20(1-54)::gfp*. The region encoding *tomm-20(1-54)::gfp* was replaced with *slc-25A46::(GGGGS)3::mCherry.* Total worm cDNA from the N2 strain was prepared as previously described ^38^. PCR primers are listed in Supplementary Table S2. *slc-25A46* cDNA was amplified by PCR using KOD FX neo DNA polymerase (TOYOBO, Tokyo Japan) with worm cDNA as the template. The slc-25A46_F_NheI and slc-25A46_R_KpnI were employed. The resulting fragment was replaced with *tomm-20(1-54)* using NheI and KpnI enzymes. Subsequently, the *gfp* sequence was replaced with *(GGGGS)3::mCherry* using Gibson assembly. *(GGGGS)3* was inserted as a flexible linker sequence^39^. To generate the plasmid encoding *flp-15p::fzo-1::(GGGGS)3::mCherry,* the *fzo-1* sequence was amplified from extracted N2 genome DNA by PCR using KOD Plus high-fidelity DNA polymerase (TOYOBO, Tokyo Japan). The region encoding *slc-25A46* was replaced with the *fzo-1* sequence.

### Worm experiments

Worm strains are described in Supplementary Table S3. All *C. elegans* strains were cultured in standard nematode growth medium (NGM) plates seeded with OP50 *Escherichia coli* at 20 ^32^. The wild-type strain, N2 Bristol, and SD1347 carrying ccIs4251, were obtained from *C. elegans* genetic center (Minneapolis, MN, USA). The deletion mutant *fzo-1(tm1133*), *eat-3(tm1107)*, and *drp-1(tm1108)* were obtained from National Bioresource Project of Japan (Tokyo Women’s Medical University School of Medicine, Japan). The PVD neuron marker *wyIs592* [*ser-2prom3p::myr-gfp*] is previously described ^36^. For transformation of worms, DNA injection was performed as described ^40^. Plasmids encoding *flp-15p::tomm-20(1-54aa)::gfp* and *odr-1p::gfp* were injected to establish *jpnEx15*[*flp-15p::tomm-20(1-54aa)::gfp*, *flp-15p::myrTagRFP-T, odr-1p::gfp*]. The extrachromosomal array was integrated to the genome by UV irradiation. The resultant integration was named *jpnIs4*. For *slc-25A46::mCherry* and *fzo-1::mCherry* transgenic worms, *flp-15p::slc-25A46::(GGGGS)3::mCherry* (20 ng/µl) or *flp-15p::fzo-1::(GGGGS)3::mCherry* (20 ng/µl) plasmid was injected into *jpnIs4* gonads, respectively. pCFJ90 (2.5 ng/µl) and pBlueScript II KS(-) (80 ng/µl) were used as co-injection markers. Then, worms carrying the extrachromosomal array were crossed to each mitochondrial mutant.

### EMS mutagenesis screening

EMS mutagenesis was performed as described previously ^32^ ^31^. The worm carrying *jpnIs4* was crossed with N2 for 8 times and used as a starting strain. For suppressor screens, *slc-25A46(jpn15); jpnIs4* was used as a starting strain. Synchronized L4 hermaphrodites were mutagenized with 0.1 M EMS (ethyl methanesulfonate, #M0880-1G, Sigma-Aldrich). Mutagenized worms were picked onto fresh plates and let them lay eggs. F2 worms were screened under a fluorescent microscopy, and candidates were identified by the abnormal mitochondrial morphology and localization in PHA neurons. The candidates were backcrossed to the parent line three times. Mutant locus was determined by single-nucleotide polymorphism (SNP) mapping^41^. For *jpn15*, whole genome sequencing was outsourced to Eurofins genomics Japan (Tokyo, Japan). Genomic data was analyzed using the Galaxy platform as described previously ^42^ ^38^. Reference genome sequence (WBcel235.75.fasta) and gene annotations (WBcel235.75.gtf) were obtained from WormBase. All of the following sequence analysis were performed using default parameters predefined in the Galaxy platform. On the Galaxy platform, a pair of FASTQ files obtained from Eurofins were firstly trimmed with Trim Galore!. Aligning sequencing reads to the genome sequence was performed using Bowtie 2. Variant search was performed using FreeBayes. Variants were annotated to genes using Snpeff. By comparing the variant data and the SNP mapping data, we found a nonsense mutation in *slc-25A46* gene. The mutation was confirmed by genomic PCR followed by Sanger sequencing using CEQ8000 (Beckman Coulter Inc, Brea, CA, USA).

For *jpn73*, SNP mapping indicated that the genomic region, which includes *drp-1*, contained the causative mutation. Therefore, the genomic region of the *drp-1* gene was amplified by PCR from *jpn73* mutant using drp-1 genome PCR_F (5’-GGCGTTCACAGTCAATCGAAGG-3’) and drp-1 genome PCR_R (5’-GGGAACGGAGCATAGAGATCATACAG-3’) primers. The genomic DNA fragment was separated by electrophoresis using a 1% agarose gel and purified using QIAquick Gel Extraction Kit (QIAGEN). The genome sequence of *drp-1* was determined by Sanger sequencing using primers listed in Supplementary Table S2.

### Genome editing

Co-CRISPR method was used for genome editing of worms^43^. The single-strand oligo DNA (ssODN) with *dpy-10 (cn64)* was used as a co-injection marker. Target sequences for guide RNA were inserted to an expression vector pTK73^44^. We chose two Cas9 target cites in *slc-25A46* sequence for each experiment.

To correct the *jpn15* mutation, target sites were *jpn15*_rescue_01, 5’-GAATTCAGACATTCTAGAA-3’ *jpn15*_rescue_02, 5’-CATTCTAGAAAGGAGCAAT-3’

To introduce disease associated mutations, target sites were *slc-25A46*_disease_01, 5’-GATGAACAATTGTTTCAAA-3’ *slc-25A46*_disease_02, 5’-TTCATCGAATGTATATTCA-3’

Synthesized ssODN (Eurofins genetics Japan) were used as repair templates (Supplementary Table S3). We introduced restriction enzyme cite as synonymous mutations to guide RNA recognition cites to prohibit the cleavage of the repair templates and for following genotyping process. All materials were mixed so that each final concentration became 50 ng/µl, and injected the mixture into worms. *dpy* or *rol* F1 worms were singled under a stereo microscope (Stemi 508, Carl Zeiss). Recombination was screened by PCR followed by digestion with each restriction enzyme, and then confirmed by Sanger sequencing. Obtained lines were crossed with N2 worms three times. All of the plasmids, primers and ssODNs are listed in Supplementary Table S1 and S2.

#### Fluorescent microscopy

Fluorescent signals in living worms are observed without fixation, following the previously described procedure ^45^. Worms are mounted on a 0.25mM levamisole (Sigma) that is placed on a 3% agarose pad. We used a Zeiss Axio Observer microscope equipped with an Objective C-Apochromat 40x (NA 1.2) and an LSM800 confocal microscope system (Carl Zeiss). The system was controlled using ZEN software (Carl Zeiss). Airy unit settings were configured to 2, and the Z stack mode was employed to capture mages. Parameters for resolution in the X-Y direction and thickness in the Z direction were the optimal values indicated by the ZEN software. Z-projection was performed using the same software. Mitochondrial size was measured using the ZEN software. Because mitochondria within dendrites have a thickness in the Z-direction less than the wavelength of light, mitochondrial size was expressed as an area on the Z-projection image rather than volume.

#### Rapid cooling procedure and quick freeze substitution for electron microscopy

The cryofixation of worms was conducted using the ultra-rapid cryo-technique as described previously ^46^. Briefly, adult animals were picked onto four NGM plates (8-10 animals/plate) with OP50 bacteria and cultured at 20 °C. On the third day of culture, a mixture of L1-L4 animals was collected using a cryoprotective solution (10% ethylene glycol in deionized distilled water) and incubated for more than 30 minutes at room temperature. The drop of animals was loaded on small-cut cellulose acetate membrane (Membrane filter C020A047A, ADVANTEC). After the liquid volume reduction by absorption into the membrane, the samples were dipped manually into liquid ethane or liquid propane and then transferred into liquid nitrogen. The samples were further transferred to cryotubes (Cryovial, 2 ml, TOHO) containing 2% OsO4 and 5% DDW in acetone at -80 °C. The specimens were freeze substituted by the quick freeze substitution (QFS) method ^47^, washed five times with acetone and three times with propylene oxide (PO) and infiltrated with Epon resin/PO in steps of 33% 1 h, 50% 3 h, and 100% resin overnight. After infiltration, the specimens were embedded in freshly prepared Epon resin and polymerized for 2 days at 60°C.

#### Electron microscopy

70 nm ultrathin sections were prepared onto Formvar-coated copper slot grids using an ultramicrotome (EM UC7, Leica). The sections were post-stained for 5 min in 5% uranyl acetate in water at 60°C and followed by 5 min in lead citrate. Mitochondria in body wall muscles were photographed using a JEM-2100F electron microscope (JEOL) equipped with TemCam-F216 (TVIPS) at 200 kV and 10,000×.

#### Statistical analyses and graph preparation

Statistical analyses were performed using Graph Pad Prism version 10.1.1. Statistical methods and sample size are described in the figure legends. Graphs were prepared using Graph Pad Prism version 9, exported in the EPS file format and aligned by Adobe Illustrator 2023.

## Reference

1. Giacomello, M., Pyakurel, A., Glytsou, C., and Scorrano, L. (2020). The cell biology of mitochondrial membrane dynamics. Nat Rev Mol Cell Biol 21, 204–224. 10.1038/s41580-020-0210-7.

2. Chappie, J.S., Acharya, S., Liu, Y.W., Leonard, M., Pucadyil, T.J., and Schmid, S.L. (2009). An intramolecular signaling element that modulates dynamin function in vitro and in vivo. Mol Biol Cell 20, 3561–3571. 10.1091/mbc.e09-04-0318.

3. Antonny, B., Burd, C., De Camilli, P., Chen, E., Daumke, O., Faelber, K., Ford, M., Frolov, V.A., Frost, A., Hinshaw, J.E., et al. (2016). Membrane fission by dynamin: what we know and what we need to know. EMBO J 35, 2270–2284. 10.15252/embj.201694613.

4. Alexander, C., Votruba, M., Pesch, U.E., Thiselton, D.L., Mayer, S., Moore, A., Rodriguez, M., Kellner, U., Leo-Kottler, B., Auburger, G., et al. (2000). OPA1, encoding a dynamin-related GTPase, is mutated in autosomal dominant optic atrophy linked to chromosome 3q28. Nat Genet 26, 211–215. 10.1038/79944.

5. Zuchner, S., Mersiyanova, I.V., Muglia, M., Bissar-Tadmouri, N., Rochelle, J., Dadali, E.L., Zappia, M., Nelis, E., Patitucci, A., Senderek, J., et al. (2004). Mutations in the mitochondrial GTPase mitofusin 2 cause Charcot-Marie-Tooth neuropathy type 2A. Nat Genet 36, 449–451. 10.1038/ng1341.

6. Westermann, B. (2008). Molecular machinery of mitochondrial fusion and fission. J Biol Chem 283, 13501–13505. 10.1074/jbc.R800011200.

7. Sesaki, H., and Jensen, R.E. (2001). UGO1 encodes an outer membrane protein required for mitochondrial fusion. J Cell Biol 152, 1123–1134. 10.1083/jcb.152.6.1123.

8. Sesaki, H., and Jensen, R.E. (2004). Ugo1p links the Fzo1p and Mgm1p GTPases for mitochondrial fusion. J Biol Chem 279, 28298–28303. 10.1074/jbc.M401363200.

9. Abrams, A.J., Hufnagel, R.B., Rebelo, A., Zanna, C., Patel, N., Gonzalez, M.A., Campeanu, I.J., Griffin, L.B., Groenewald, S., Strickland, A.V., et al. (2015). Mutations in SLC25A46, encoding a UGO1-like protein, cause an optic atrophy spectrum disorder. Nat Genet 47, 926–932. 10.1038/ng.3354.

10. Janer, A., Prudent, J., Paupe, V., Fahiminiya, S., Majewski, J., Sgarioto, N., Des Rosiers, C., Forest, A., Lin, Z.Y., Gingras, A.C., et al. (2016). SLC25A46 is required for mitochondrial lipid homeostasis and cristae maintenance and is responsible for Leigh syndrome. EMBO Mol Med 8, 1019–1038. 10.15252/emmm.201506159.

11. Steffen, J., Vashisht, A.A., Wan, J., Jen, J.C., Claypool, S.M., Wohlschlegel, J.A., and Koehler, C.M. (2017). Rapid degradation of mutant SLC25A46 by the ubiquitin-proteasome system results in MFN1/2-mediated hyperfusion of mitochondria. Mol Biol Cell 28, 600–612. 10.1091/mbc.E16-07-0545.

12. Wan, J., Steffen, J., Yourshaw, M., Mamsa, H., Andersen, E., Rudnik-Schoneborn, S., Pope, K., Howell, K.B., McLean, C.A., Kornberg, A.J., et al. (2016). Loss of function of SLC25A46 causes lethal congenital pontocerebellar hypoplasia. Brain 139, 2877–2890. 10.1093/brain/aww212.

13. Ali, M.S., Suda, K., Kowada, R., Ueoka, I., Yoshida, H., and Yamaguchi, M. (2020). Neuron-specific knockdown of solute carrier protein SLC25A46a induces locomotive defects, an abnormal neuron terminal morphology, learning disability, and shortened lifespan. IBRO Rep 8, 65–75. 10.1016/j.ibror.2020.02.001.

14. Suda, K., Ueoka, I., Azuma, Y., Muraoka, Y., Yoshida, H., and Yamaguchi, M. (2018). Novel Drosophila model for mitochondrial diseases by targeting of a solute carrier protein SLC25A46. Brain Res 1689, 30–44. 10.1016/j.brainres.2018.03.028.

15. Li, Z., Peng, Y., Hufnagel, R.B., Hu, Y.C., Zhao, C., Queme, L.F., Khuchua, Z., Driver, A.M., Dong, F., Lu, Q.R., et al. (2017). Loss of SLC25A46 causes neurodegeneration by affecting mitochondrial dynamics and energy production in mice. Hum Mol Genet 26, 3776–3791. 10.1093/hmg/ddx262.

16. Schuettpelz, J., Janer, A., Antonicka, H., and Shoubridge, E.A. (2023). The role of the mitochondrial outer membrane protein SLC25A46 in mitochondrial fission and fusion. Life Sci Alliance 6. 10.26508/lsa.202301914.

17. Duchesne, A., Vaiman, A., Castille, J., Beauvallet, C., Gaignard, P., Floriot, S., Rodriguez, S., Vilotte, M., Boulanger, L., Passet, B., et al. (2017). Bovine and murine models highlight novel roles for SLC25A46 in mitochondrial dynamics and metabolism, with implications for human and animal health. PLoS Genet 13, e1006597. 10.1371/journal.pgen.1006597.

18. Nguyen, M., Boesten, I., Hellebrekers, D.M., Mulder-den Hartog, N.M., de Coo, I.F., Smeets, H.J., and Gerards, M. (2017). Novel pathogenic SLC25A46 splice-site mutation causes an optic atrophy spectrum disorder. Clin Genet 91, 121–125. 10.1111/cge.12774.

19. Ichishita, R., Tanaka, K., Sugiura, Y., Sayano, T., Mihara, K., and Oka, T. (2008). An RNAi screen for mitochondrial proteins required to maintain the morphology of the organelle in Caenorhabditis elegans. J Biochem 143, 449–454. 10.1093/jb/mvm245.

20. Ren, X., Zhou, H., Sun, Y., Fu, H., Ran, Y., Yang, B., Yang, F., Bjorklund, M., and Xu, S. (2023). MIRO-1 interacts with VDAC-1 to regulate mitochondrial membrane potential in Caenorhabditis elegans. EMBO Rep 24, e56297. 10.15252/embr.202256297.

21. Zhao, Y., Song, E., Wang, W., Hsieh, C.H., Wang, X., Feng, W., Wang, X., and Shen, K. (2021). Metaxins are core components of mitochondrial transport adaptor complexes. Nat Commun 12, 83. 10.1038/s41467-020-20346-2.

22. Niwa, S., Lipton, D.M., Morikawa, M., Zhao, C., Hirokawa, N., Lu, H., and Shen, K. (2016). Autoinhibition of a Neuronal Kinesin UNC-104/KIF1A Regulates the Size and Density of Synapses. Cell Rep 16, 2129–2141. 10.1016/j.celrep.2016.07.043.

23. Meyerzon, M., Fridolfsson, H.N., Ly, N., McNally, F.J., and Starr, D.A. (2009). UNC-83 is a nuclear-specific cargo adaptor for kinesin-1-mediated nuclear migration. Development 136, 2725–2733. 10.1242/dev.038596.

24. Campbell, D., and Zuryn, S. (2024). The mechanisms and roles of mitochondrial dynamics in C. elegans. Semin Cell Dev Biol 156, 266–275. 10.1016/j.semcdb.2023.10.006.

25. Rolland, S.G., Lu, Y., David, C.N., and Conradt, B. (2009). The BCL-2-like protein CED-9 of C. elegans promotes FZO-1/Mfn1,2- and EAT-3/Opa1-dependent mitochondrial fusion. J Cell Biol 186, 525–540. 10.1083/jcb.200905070.

26. Scholtes, C., Bellemin, S., Martin, E., Carre-Pierrat, M., Mollereau, B., Gieseler, K., and Walter, L. (2018). DRP-1-mediated apoptosis induces muscle degeneration in dystrophin mutants. Sci Rep 8, 7354. 10.1038/s41598-018-25727-8.

27. Labrousse, A.M., Zappaterra, M.D., Rube, D.A., and van der Bliek, A.M. (1999). C. elegans dynamin-related protein DRP-1 controls severing of the mitochondrial outer membrane. Mol Cell 4, 815–826. 10.1016/s1097-2765(00)80391-3.

28. Reese, T.S. (1965). Olfactory Cilia in the Frog. J Cell Biol 25, 209–230. 10.1083/jcb.25.2.209.

29. Kanaji, S., Iwahashi, J., Kida, Y., Sakaguchi, M., and Mihara, K. (2000). Characterization of the signal that directs Tom20 to the mitochondrial outer membrane. J Cell Biol 151, 277–288. 10.1083/jcb.151.2.277.

30. Inglis PN., O.G., Lerouz MR., Scholey JM. (2006). The sensory cilia of Caenorhabditis elegans (November 27, 2006). WormBook.

31. Niwa, S. (2016). The nephronophthisis-related gene ift-139 is required for ciliogenesis in Caenorhabditis elegans. Sci Rep 6, 31544. 10.1038/srep31544.

32. Brenner, S. (1974). The genetics of Caenorhabditis elegans. Genetics 77, 71–94. 10.1093/genetics/77.1.71.

33. Thompson, O., Edgley, M., Strasbourger, P., Flibotte, S., Ewing, B., Adair, R., Au, V., Chaudhry, I., Fernando, L., Hutter, H., et al. (2013). The million mutation project: a new approach to genetics in Caenorhabditis elegans. Genome Res 23, 1749–1762. 10.1101/gr.157651.113.

34. Oren-Suissa, M., Gattegno, T., Kravtsov, V., and Podbilewicz, B. (2017). Extrinsic Repair of Injured Dendrites as a Paradigm for Regeneration by Fusion in Caenorhabditis elegans. Genetics 206, 215–230. 10.1534/genetics.116.196386.

35. Brar, H.K., Dey, S., Bhardwaj, S., Pande, D., Singh, P., Dey, S., and Ghosh-Roy, A. (2022). Dendrite regeneration in C. elegans is controlled by the RAC GTPase CED-10 and the RhoGEF TIAM-1. PLoS Genet 18, e1010127. 10.1371/journal.pgen.1010127.

36. Dong, X., Liu, O.W., Howell, A.S., and Shen, K. (2013). An extracellular adhesion molecule complex patterns dendritic branching and morphogenesis. Cell 155, 296–307. 10.1016/j.cell.2013.08.059.

37. Nakano, J., Chiba, K., and Niwa, S. (2022). An ALS-associated KIF5A mutant forms oligomers and aggregates and induces neuronal toxicity. Genes Cells 27, 421–435. 10.1111/gtc.12936.

38. Higashida, M., and Niwa, S. (2023). Dynein intermediate chains DYCI-1 and WDR-60 have specific functions in Caenorhabditis elegans. Genes Cells 28, 97–110. 10.1111/gtc.12996.

39. Trinh, R., Gurbaxani, B., Morrison, S.L., and Seyfzadeh, M. (2004). Optimization of codon pair use within the (GGGGS)3 linker sequence results in enhanced protein expression. Mol Immunol 40, 717–722. 10.1016/j.molimm.2003.08.006.

40. Mello, C.C., Kramer, J.M., Stinchcomb, D., and Ambros, V. (1991). Efficient gene transfer in C.elegans: extrachromosomal maintenance and integration of transforming sequences. EMBO J 10, 3959–3970. 10.1002/j.1460-2075.1991.tb04966.x.

41. Davis, M.W., Hammarlund, M., Harrach, T., Hullett, P., Olsen, S., and Jorgensen, E.M. (2005). Rapid single nucleotide polymorphism mapping in C. elegans. BMC Genomics 6, 118. 10.1186/1471-2164-6-118.

42. Community, G. (2022). The Galaxy platform for accessible, reproducible and collaborative biomedical analyses: 2022 update. Nucleic Acids Res 50, W345–W351. 10.1093/nar/gkac247.

43. Arribere, J.A., Bell, R.T., Fu, B.X., Artiles, K.L., Hartman, P.S., and Fire, A.Z. (2014). Efficient marker-free recovery of custom genetic modifications with CRISPR/Cas9 in Caenorhabditis elegans. Genetics 198, 837–846. 10.1534/genetics.114.169730.

44. Obinata, H., Sugimoto, A., and Niwa, S. (2018). Streptothricin acetyl transferase 2 (Sat2): A dominant selection marker for Caenorhabditis elegans genome editing. PLoS One 13, e0197128. 10.1371/journal.pone.0197128.

45. Anazawa, Y., and Niwa, S. (2022). Analyzing the Impact of Gene Mutations on Axonal Transport in Caenorhabditis Elegans. Methods Mol Biol 2431, 465–479. 10.1007/978-1-0716-1990-2_25.

46. Irdani, T., Fortunato, A., and Torre, R. (2015). An ultra-rapid cryo-technique for complex organisms. Cryobiology 71, 391–397. 10.1016/j.cryobiol.2015.10.144.

47. McDonald, K.L., and Webb, R.I. (2011). Freeze substitution in 3 hours or less. J Microsc 243, 227–233. 10.1111/j.1365-2818.2011.03526.x.

