## Supplemental files for "SLC-25A46 Regulates Mitochondrial Fusion through FZO-1/Mitofusin and is Essential for Maintaining Neuronal Morphology"

Figure S1

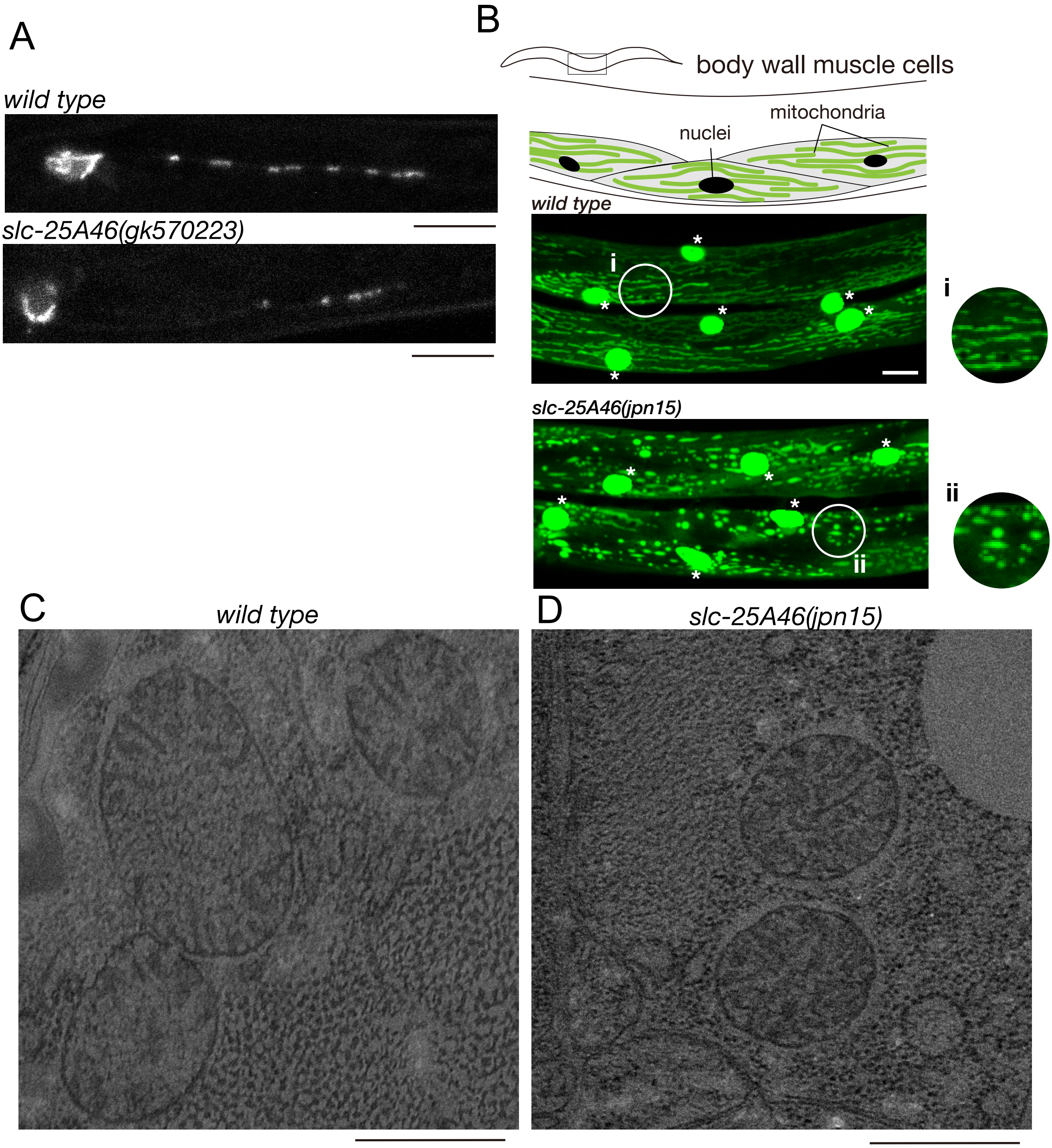


**Figure S1**
(A) Representative images showing the morphology and distribution of mitochondria in PHA neuron of wild type and slc-25A46(gk570223). Bars, 10 μm.
(B) Representative fluorescent microscopic images showing the morphology of mitochondria in muscular cells of wild type and slc-25A46(jpn15). Bars, 10 μm.
(C and D) Representative transmission electron microscopic images showing the morphology of mitochondria in muscular cells of wild type and slc-25A46(jpn15). Bars, 100 nm.

Figure S2

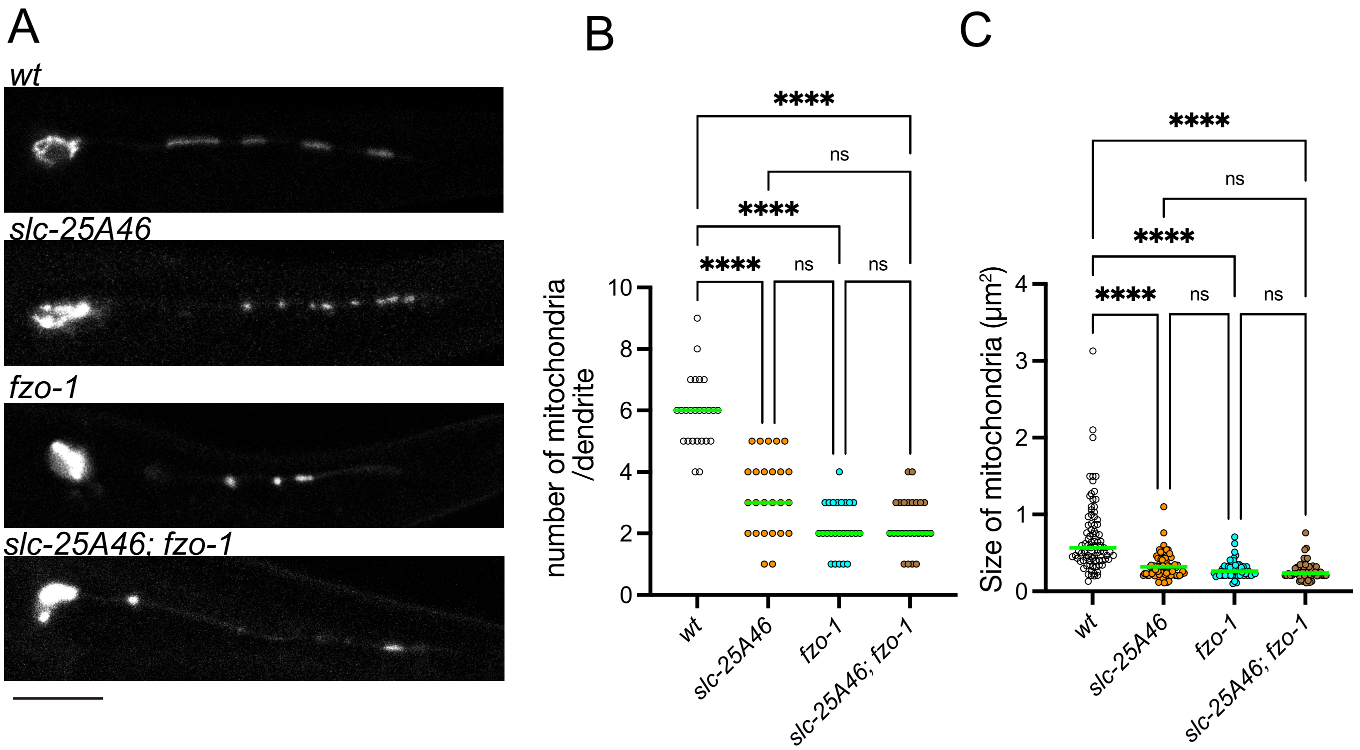


**Figure S2**

(A) Representative images showing the morphology and distribution of mitochondria in PHA neuron of wild type, slc-25A46(jpn15), fzo-1 and slc-25A46(jpn15); fzo-1. Bar, 10 μm.

(B) Dot plots showing the number of mitochondria in the PHA dendrite. Each dot shows the number of mitochondria in a single PHA dendrite. Green bars represent median values. n = 20 dendrites from 20 worms. Kruskal-Wallis test followed by Dunn's multiple comparison test. ***, $p < 0.001$. ****, $p < 0.0001$. ns, $p > 0.05$ and statistically not significant.

(C) Dot plots showing the size distribution of mitochondria in the PHA dendrite. Each dot represents the size of an individual mitochondrion in the PHA dendrite. Green bars represent median values. n = 85, 83 and 85 mitochondria. Kruskal-Wallis test followed by Dunn's multiple comparison test. ***, $p < 0.001$. ****, $p < 0.0001$. ns, $p > 0.05$ and statistically not significant.

Table S1

| plasmid name | insert | description |
|---|---|---|
| Podr-1::gfp | *odr-1p::gfp* | from Kang Shen lab (Stanford Univ.) |
| tomm-20(1-54aa)::gfp | *tomm-20(1-54aa)::gfp* | from Kang Shen lab (Stanford Univ.) |
| Pflp-15::gfp | *flp-15p::gfp* | Niwa(2015) Scientific reports |
| pSN290 | *flp-15p::tomm-20(1-54aa)::gfp* | This study |
| pSN291 | *flp-15p::myrTagRFP-T* | This study |
| pCFJ90 | *myo-2p::mCherry::unc-54utr* | from Addgene |
| pOB28 | *flp-15p::slc-25A46(cDNA)::(GGGGS)3::mCherry::unc-54_3'UTR* | This study |
| pOB29 | *flp-15p::fzo-1 genomic DNA::(GGGGS)3::mCherry::unc-54_3'UTR* | This study |
| pTK73 | *CeU6 promoter ::sgRNA (F+E) scaffold sequence* | This study |
| pTK73_rescue-1 | *CeU6 promoter :: target sequence (GAATTCAGACATTCTAGAA) :: sgRNA (F+E) scaffold sequence* | This study |
| pTK73_rescue-2 | *CeU6 promoter :: target sequence (CATTCTAGAAAGGAGCAAT) :: sgRNA (F+E) scaffold sequence* | This study |
| pTK73_P299L&E301D&R306C-1 | *CeU6 promoter :: target sequence (GATGAACAATTGTTTCAAA) :: sgRNA (F+E) scaffold sequence* | This study |
| pTK73_P299L&E301D&R306C-2 | *CeU6 promoter :: target sequence (TTCATCGAATGTATATTCA) :: sgRNA (F+E) scaffold sequence* | This study |

**Table S1 Plasmid list**

Table S2

| oligo name | sequence | comment |
|---|---|---|
| **slc-25A46_F_NheI** | atGCTAGC ATGCCTACACAATTCATTAGGAACCGG | cloning of slc-25A46 |
| **slc-25A46_R_KpnI** | atgc GGTACC cc ACCCGAAAATGGGTCTCCAGACGAC | cloning of slc-25A46 |
| **drp-1 genome PCR_F** | GGCGTTCACAGTCAATCGAAGG | sequencing of drp-1 |
| **drp-1 genome PCR_R** | GGGAACGGAGCATAGAGATCATACAG | sequencing of drp-1 |
| **jpn73_seq_primer_exon1** | TTCGCACGGCATCGAAGTCTGG | sequencing of drp-1 |
| **drp-1exon2-1_seqF** | TACTGGCTCTAAGGTTTTCACAG | sequencing of drp-1 |
| **drp-1exon2-2_seqF** | CAGGATTTCGCTACTTCGGAGCC | sequencing of drp-1 |
| **drp-1exon3-1_seq** | TATTTGGCAAAGAGATTGAATATGG | sequencing of drp-1 |
| **drp-1exon3-2_seq** | AATGCAACGAATGGTTCAGCATTGC | sequencing of drp-1 |
| **drp-1exon4_seq** | tgccaggaagtgcggatgactg | sequencing of drp-1 |
| **drp-1exon5&6_seq** | ATGTCGCTATTATCGgtatgagacc | sequencing of drp-1 |
| **jpn15jpn33_PCR_F** | CTGAGCCTCACCCAATCTCGAAATC | sequencing of jpn15 and jpn33 |
| **jpn15jpn33_PCR_R** | ATTCGATCCACACCCTTCACAAGAAC | sequencing of jpn15 and jpn33 |
| **jpn15jpn33_seq_primer** | TCGCTAATTTCCCATCCATGCGGTG | sequencing of jpn15 and jpn33 |
| **dpy-10(cn64)_oligo** | TGAAGCCATGTGAAGCTCCGCTACCATAGGCACCACAAGCGGTACGG GTTCCAGTCATTCTCATCTTGCCGTATTGAAGTTCAAGTGCAGCCTCG TCGTTTGATCTC | repair template for dpy-10 |
| **P299L_ssODN** | ATCATCAATGGTGTTAACTGATTTGATACTTTATCTTTTCGAGACCATC GTGCACAGAATGTACATCCAAGGAACACGAACACTTATTGATAATAT GGAT | repair template for disease mutation |
| **E301D_ssODN** | CATCAATGGTGTTAACTGATTTGATACTTTATCCATTCGATACCATCGT GCACAGAATGTACATCCAAGGAACACGAACACTTATTGATAATATGGATAC | repair template for disease mutation |
| **slc_R306C_ssODN** | ATCAATGGTGTTAACTGATTTGATACTTTATCCATTCGAGACCATCGTGC ACTGCATGTACATCCAAGGAACACGAACACTTATTGATAATATGGATACA | repair template for disease mutation |
| **jpn15rescue_ssODN** | CTGATTCCTGTAATCTGTAATTCTGTCGCTAAACAAGGAATCCAAACC TTTTGGAAGGGCGCCATCGGCTCAAGTGTGCTCTGGGGCCTCACGAA TGTTACGGAA | introducing jpn33 mutation in jpn15 |

**Table S2 List of oligo DNAs used in this study**

Table S3

| strain name | genotype | source and commments |
|---|---|---|
| N2 | *wild type* | from CGC |
| OTL45 | *jpnEx15[flp-15p::mito::GFP; flp-15p::myrTagRFP-T; odr-1p::GFP]* | this study, mitochondria is visualized with GFP in the PHA neuron |
| OTL48 | *jpnIs4[flp-15p::mito::GFP; flp-15p::myrTagRFP-T; odr-1p::GFP] V* | this study, mitochondria is visualized with GFP in the PHA neuron |
| OTL247 | *slc-25A46(jpn15)I; jpnIs4V* | this study, null mutant of slc-25A46 |
| OTL249 | *slc-25A46(jpn33)I; jpnIs4V* | this study, *jpn15* mutation was corrected by CRISPR/Cas9 |
| VC40317 | *slc-25A46(gk570223)* | from CGC |
| OTL277 | *slc-25A46(gk570223); jpnIs4V* | this study |
| | *fzo-1(tm1133)* | from NBRP |
| | *eat-3(tm1107)* | from NBRP |
| | *drp-1(tm1108)* | from NBRP |
| OTL251 | *fzo-1(tm1133); jpnIs4V* | this study |
| OTL250 | *eat-3(tm1107); jpnIs4V* | this study |
| OTL252 | *drp-1(tm1108); jpnIs4V* | this study |
| OTL270 | *jpnIs4 ; jpnEx53[Posm-6::slc-25A46,myo-2p::RFP ]* | this study |
| OTL275 | *fzo-1; jpnIs4 ; jpnEx53[Posm-6::slc-25A46, myo-2p::RFP ]* | this study |
| OTL271 | *slc-25A46; jpnIs4 ; jpnEx53[Posm-6::slc-25A46,myo-2p::RFP ]* | this study |
| OTL276 | *jpnIs4 ; jpnEx52[Posm-6::slc-25A46,myo-2p::RFP]* | this study |
| OTL241 | *slc-25A46(jpn15); drp-1(jpn73); jpnIs4* | this study |
| OTL242 | *drp-1(jpn73); jpnIs4V* | this study |
| OTL244 | *jpnIs4V; jpnEx104 [flp-15p::fzo-1, myo-2p::RFP]* | this study |
| OTL245 | *slc-25A46(jpn15); jpnIs4V; jpnEx104 [flp-15p::fzo-1, myo-2p::RFP]* | this study |
| OLT246 | *fzo-1(tm1133);jpnIs4; jpnEx104 [flp-15p::fzo-1, myo-2p::RFP]* | this study |
| OTL253 | *slc-25A46(jpn37); jpnIs4V* | this study, P299L mutation in the slc-25A46 gene |
| OTL254 | *slc-25A46(jpn38); jpnIs4V* | this study, E301D mutation in the slc-25A46 gene |
| OTL255 | *slc-25A46(jpn39); jpnIs4V* | this study, R306C mutation in the slc-25A46 gene |
| TV15911 | *wyIs592 [ser-2prom-3p::myr-GFP ]* | from Kang Shen lab(Stanford Univ.), GFP is expressed in the PVD neuron |
| OTL259 | *slc-25A46(jpn15); wyIs592* | this study |
| OTL256 | *slc-25A46(jpn37); wyIs592* | this study, P299L mutation in the slc-25A46 gene |
| OTL257 | *slc-25A46(jpn38); wyIs592* | this study, E301D mutation in the slc-25A46 gene |
| OTL58 | *slc-25A46(jpn39); wyIs592* | this study, R306C mutation in the slc-25A46 gene |
| SD1347 | *ccIs4251* | from CGC |
| OTL260 | *slc-25A46(jpn15);ccIs4251* | this study |

**Table S3 Strain list**
